# Human demographic history has amplified the effects of background selection across the genome

**DOI:** 10.1101/181859

**Authors:** Raul Torres, Zachary A. Szpiech, Ryan D. Hernandez

## Abstract

Natural populations often grow, shrink, and migrate over time. Demographic processes such as these can impact genome-wide levels of genetic diversity. In addition, genetic variation in functional regions of the genome can be altered by natural selection, which drives adaptive mutations to higher frequencies or purges deleterious ones. Such selective processes impact not only the sites directly under selection but also nearby neutral variation through genetic linkage through processes referred to as genetic hitch-hiking in the context of positive selection and background selection (BGS) in the context of purifying selection. While there is extensive literature examining the impact of selection at linked sites at demographic equilibrium, less is known about how non-equilibrium demographic processes impact the effects of hitchhiking and BGS. Utilizing a global sample of human whole-genome sequences from the Thousand Genomes Project and extensive simulations, we investigate how non-equilibrium demographic processes magnify and dampen the consequences of selection at linked sites across the human genome. When binning the genome by inferred strength of BGS, we observe that, compared to Africans, non-African populations have experienced larger proportional decreases in neutral genetic diversity in such regions. We replicate these findings in admixed populations by showing that non-African ancestral components of the genome have also been impacted more severely in these regions. We attribute these differences to the strong, sustained/recurrent population bottlenecks that non-Africans experienced as they migrated out of Africa and throughout the globe. Furthermore, we observe a strong correlation between *F*_ST_ and inferred strength of BGS, suggesting a stronger rate of genetic drift. Forward simulations of human demographic history with a model of BGS support these observations. Our results show that non-equilibrium demography significantly alters the consequences selection at linked sites and support the need for more work investigating the dynamic process of multiple evolutionary forces operating in concert.

**Author summary:** Patterns of genetic diversity within a species are affected at broad and fine scales by population size changes (“demography”) and natural selection. From both population genetics theory and observation of genomic sequence data, it is known that demography can alter genome-wide average neutral genetic diversity. Additionally, natural selection can affect neutral genetic diversity regionally across the genome via selection at linked sites. During this process, natural selection acting on adaptive or deleterious variants in the genome will also impact diversity at nearby neutral sites due to genetic linkage. However, less is well known about the dynamic changes to diversity that occur in regions impacted by selection at linked sites when a population undergoes a size change. We characterize these dynamic changes using thousands of human genomes and find that the population size changes experienced by humans have shaped the consequences of linked selection across the genome. In particular, population contractions, such as those experienced by non-Africans, have disproportionately decreased neutral diversity in regions of the genome inferred to be under strong background selection (i.e., selection at linked sites that is caused by natural selection acting on deleterious variants), resulting in large differences between African and non-African populations.

## Introduction

Genetic diversity in a species is determined through the complex interplay of mutation, demography, genetic drift, and natural selection. These evolutionary forces operate in concert to shape patterns of diversity at both the local scale and genome-wide scale. For example, in recombining species, levels of genetic diversity are distributed heterogeneously across the genome as peaks and valleys that are often correlated with recombination rate and generated by past or ongoing events of natural selection [1]. But at the genome-wide scale, average levels of genetic diversity are primarily impacted by population size changes, yielding patterns of diversity that are a function of a population’s demographic history [2]. These patterns of diversity may also yield information for inferring past events of natural selection and population history, giving valuable insight into how populations have evolved over time [3–8]. With recent advances in sequencing technology yielding whole-genome data from thousands of individuals from species with complex evolutionary histories [9,10], formal inquiry into the interplay of demography and natural selection and testing whether demographic effects act uniformly across the genome as a function of natural selection is now possible.

In the past decade, population genetic studies have shed light on the pervasiveness of dynamic population histories in shaping overall levels of genetic diversity across different biological species. For example, multiple populations have experienced major population bottlenecks and founder events that have resulted in decreased levels of genome-wide diversity. Evidence for population bottlenecks exists in domesticated species such as cattle [11], dogs [12], and rice [13], and in natural populations such as *Drosoph ila melanogaster* [14–16], rhesus macaque [17], and humans [18,19]. Notably, population bottlenecks leave long lasting signatures of decreased diversity, which may be depressed even after a population has recovered to or surpassed its ancestral size [20,21]. Such examples are evident in humans, where non-African populations exhibit a lower amount of genetic diversity compared to Africans [9], despite the fact that they have been inferred to have undergone a greater population expansion in recent times [22,23].

Locally (i.e., regionally) across the genome, the action of natural selection can also lead to distinct signatures of decreased genetic diversity (although some forms of selection, such as balancing selection, can increase genetic diversity [24]). For example, mutations with functional effects may be removed from the population due to purifying selection or fix due to positive selection, thereby resulting in the elimination of genetic diversity at the site. But while sites under direct natural selection in the genome represent only a small fraction of all sites genome-wide, the action of natural selection on these selected sites can have far-reaching effects across neutral sites in the genome due to linkage. Under positive selection, genetic hitchhiking [25] causes variants lying on the same haplotype as the selected allele to rise to high frequency during the selection process (note that we will use the term “genetic hitchhiking” here only in the positive selection context of selection at linked sites). Conversely, under purifying selection, background selection (BGS) [26] causes linked neutral variants to decrease in frequency or be removed from the population. Both of these processes of selection at linked sites result in decreased neutral genetic diversity around the selected site. Recombination can decouple neutral sites from selected sites in both cases and neutral diversity tends to increase toward its neutral expectation as genetic distance from selected sites increases [27].

Evidence for genetic hitchhiking and BGS has been obtained from the genomes of several species, including *Drosophila melanogaster* [28–33], wild and domesticated rice [34,35], nematodes [36,37], humans [3,6,38–42], and others (see [1] for a review). While the relative contributions of genetic hitchhiking and BGS to shaping patterns of human genomic diversity have been actively debated [40,43–45], the data strongly support a large role for BGS in shaping genome-wide patterns of neutral genetic variation [41,42]. Indeed, recent arguments have been made in favor of BGS being treated as the null model when investigating the impact of selection at linked sites across recombining genomes [1,32,45–48], with one study in humans showing that BGS has decreased genetic diversity by 19–26% if other modes of selection at linked sites are assumed to be minor [6].

Although the effects of selection at linked sites across the genome have been described in a multitude of studies, it is still less obvious whether populations that have experienced different demographic histories, such as African and non-African human populations, should exhibit similar relative effects in those regions. In the context of BGS, early work resulted in the expression π ≈ *4f_0_N_e_μ* [26], which suggests that the expected level of diversity with BGS is simply proportional to the neutral expectation (with proportionality constant *f*_0_ being a function of the rates of deleterious mutation and recombination). Importantly, while populations with different *N_e_* are expected to have different levels of π in the context of BGS, the proportionality constant *f_0_* is assumed to be independent of population history (so long as the recombination landscape and deleterious mutation rates remain constant). Much of the theory developed in the context of BGS has been developed under the assumption that the population is at demographic equilibrium, and recent work has demonstrated that this assumption likely holds under changing demography if selection is strong enough (or populations are large enough) such that mutation-selection balance is maintained [49,50]. However, strong, sustained population bottlenecks may lead to violations of that assumption. Thus, in humans and other natural populations experiencing non-equilibrium demography, genetic drift may perturb genetic diversity at neutral sites under BGS that are unaccounted for in these models.

While little attention has been given to the potential consequences of demography on patterns of neutral variation in regions experiencing selection at linked sites (but see [51,52] for how selection at linked sites may impact the inference of demography itself), recent studies have suggested that alleles directly under natural selection experience non-linear dynamics in the context of non-equilibrium demography. For the case of purifying selection, the equilibrium frequency of an allele is dependent on its fitness effect, with deleterious alleles having lower equilibrium frequencies than neutral alleles. After a population size change, deleterious alleles tend to change frequency faster than neutral alleles, allowing them to reach their new equilibrium frequency at a faster rate [53,54]. This can result in relative differences in deleterious allele frequencies among populations with different demographic histories. Such effects are especially apparent in populations suffering bottlenecks [55] and have been tested and observed between different human populations with founder populations exhibiting a greater proportion of non-synonymous variation relative to synonymous variation [56–58]. We hypothesized that these non-equilibrium dynamics could also perturb nearby neutral variants due to the effects of selection at linked sites. In support of our hypothesis, a recent simulation study modeling *Drosophila* observed that population bottlenecks can result in different rates of recovery of neutral genetic diversity depending on the strength of BGS [48]. Another recent study [59] analyzed neutral diversity surrounding putatively deleterious loci in domesticated versus wild maize. They found that the extreme domestication bottleneck reduced the efficiency of purifying selection, which has resulted in higher relative diversity in the domesticated population compared to the wild population (which has likely experienced a much more stable demographic history). Together, these studies provide further evidence that non-equilibrium demography should have a strong effect on patterns of diversity in the presence of selection at linked sites.

To investigate the impact of non-equilibrium dynamics in regions experiencing selection at linked sites, we measure patterns of average pairwise neutral genetic diversity (π), or neutral heterozygosity if the population was admixed, as a function of the strength of BGS, *B* (inferred by Ref. [6]), within a global set of human populations from phase 3 of the Thousand Genomes Project (TGP) [9]. We focus on the ratio of neutral diversity in regions of strong BGS (low B) to regions of weak BGS (high B, the closest proxy available for neutral variation in humans), which we term “relative diversity.” While there are many caveats that may plague the direct interpretation of specific *B* values (e.g., positive selection is not modeled, the distribution of fitness effects are inconsistent with other studies, and the deleterious mutation rate exceeds the per base pair mutation rate of other studies), we argue below that *B* scores nevertheless provide a decent proxy for ranking sites from most closely linked to deleterious loci (low B) to most unlinked from deleterious loci (high *B*) in humans.

We find substantial differences in relative diversity between populations, which we attribute to their non-equilibrium demographics. We confirm that the interplay of demography and selection at linked sites can explain the differences of relative diversity across human populations with simulations incorporating a parametric demographic model of human history [7] with and without a model of BGS. We also investigate how genetic differentiation between TGP populations (as measured by *F_ST_*) is shaped by selection at linked sites by measuring *F_ST_* as a function of *B*. Finally, we demonstrate that back migration from Europeans and Asians into Africa re-introduces sufficient deleterious variation to impact patterns of BGS, leading to decreased relative diversity in Africans. Our results demonstrate the strong impact that changing demography has on perturbing levels of diversity in regions experiencing selection at linked sites and have implications for population genetic studies seeking to characterize linked selection across any species or population that is not at demographic equilibrium.

## Results

### Differential impact of selection at linked sites across human populations

We measured mean pairwise genetic diversity (π) in the autosomes (we ignore the sex chromosomes and the mitochondrial genome for all analyses) among the 20 non-admixed populations from the phase 3 TGP data set, consisting of 5 populations each from 4 continental groups: Africa (AFR), Europe (EUR), South Asia (SASN), and East Asia (EASN; population labels and groupings reported in S12 Table in Supporting information). A set of stringent filters, including the masking of sites inferred to be under selective sweeps, were first applied to all 20 populations to identify a high-quality set of putatively neutral sites in the genome (see Materials and Methods). Sites were then divided into quantile bins based on *B* [6]. For our initial set of analyses, we focused on the bins corresponding to the 1% of sites inferred to be under the strongest amount of BGS (i.e., sites having the lowest inferred *B* values) and the 1% of sites inferred to be under the weakest amount BGS (i.e., sites having the highest inferred *B* values). Mean diversity was normalized by divergence with rhesus macaque within these bins for each population and is shown in Figs 1A-B. As expected, normalized diversity was highest in African populations and lowest in East Asian populations across both 1% *B* quantile bins.

**Fig 1.**
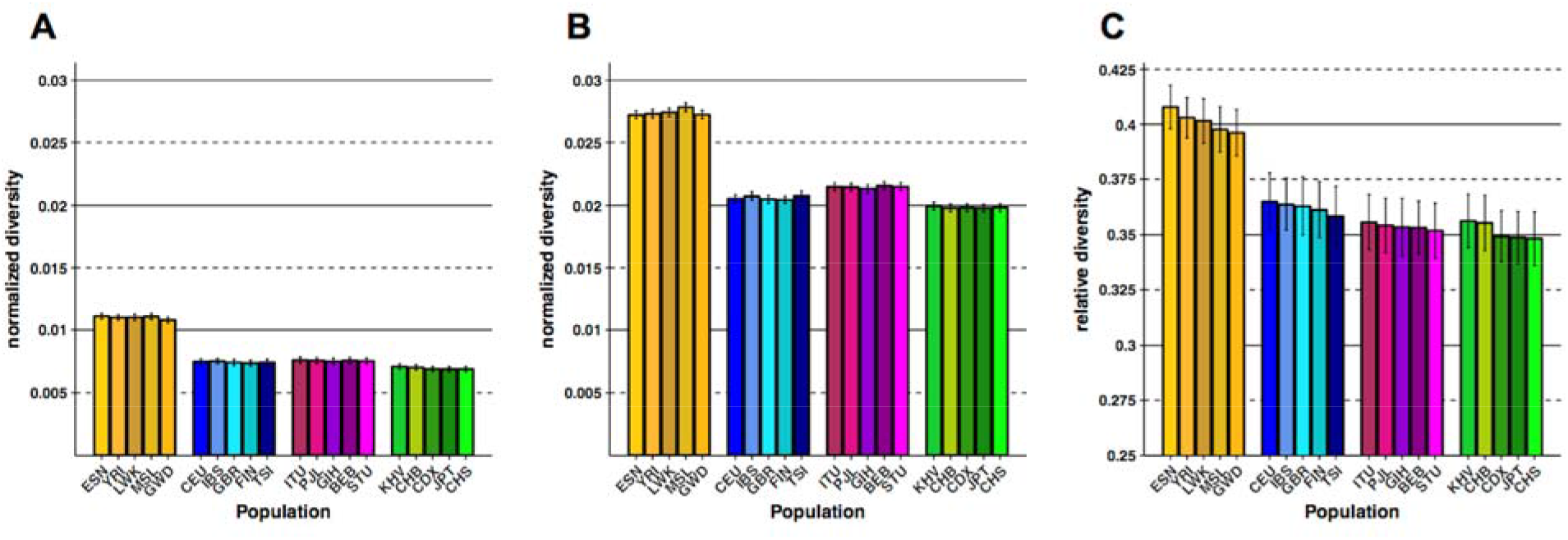
Normalized diversity and relative diversity for non-admixed populations of the Thousand Genomes Project (TGP) (A) Normalized diversity (π/divergence) measured across the lowest 1% *B* quantile bin (strong BGS). (B) Normalized diversity measured across the highest 1% *B* quantile bin (weak BGS). (C) Relative diversity: the ratio of normalized diversity in the lowest 1% *B* bin to normalized diversity in the highest 1% *B* bin (π/π_0_). TGP population labels are indicated below each bar (see S12 Table in Supporting information for population label descriptions), with African populations colored by gold shades, European populations colored by blue shades, South Asian populations colored by violet shades, and East Asian populations colored by green shades. Error bars represent ±1 SEM calculated from 1,000 bootstrapped datasets.

To estimate the impact that selection at linked sites has had on neutral diversity, we calculated a statistic called “relative diversity” (analogous to π/π_0_ in the BGS literature; [26,60]) for each population. We define relative diversity as the ratio of normalized diversity in the lowest 1% *B* bin to normalized diversity in the highest 1% *B* bin, which should capture the relative impact of selection at linked sites within the genome. Fig 1C shows that relative diversity was lower in non-African populations (0.348-0.365 for non-Africans, 0.396-0.408 for Africans), suggesting that these populations have experienced a greater reduction in diversity due to selection at linked sites than can be explained by demography alone.

To characterize these effects across a broader distribution of sites experiencing selection at linked sites, we grouped populations together according to their continental group (i.e., African, European, South Asian, and East Asian, see S12 Table in Supporting information for a detailed description) and estimated relative diversity at neutral sites for each of the continental groups in bins corresponding to the lowest 1%, 5%, 10%, and 25% quantiles of *B* (note these partitions were not disjoint). As expected, relative diversity increased for all continental groups as the bins became more inclusive (Fig 2B), reflecting a reduced impact on the reduction of diversity due to selection at linked sites. We also observed that non-African continental groups consistently had a lower relative diversity compared to African groups, demonstrating that the patterns we observed in the most extreme regions experiencing selection at linked sites also held for broader regions. Interestingly, we observed a consistent trend of rank order for relative diversity between the different continental groups for each quantile bin, with the East Asian group experiencing the greatest reduction of relative diversity, followed by the South Asian, European, and African groups. This result suggested a stronger effect for demography on the diversity-reducing effect selection at linked sites for those populations experiencing the strongest bottlenecks. However, the observed differences in relative diversity between non-African and African continental groups became less pronounced as the bins became more inclusive (Fig 2B). These effects remained even after we controlled for the effects of GC-biased gene conversion and recombination hotspots (S2 and S4 Figs) or if we did not normalize diversity by divergence (S3 and S5 Figs). Patterns of relative diversity in regions of local ancestry (i.e., African, European, or Native American) across admixed TGP populations also largely recapitulated the patterns observed in their continental group counterparts across *B* quantile bins, with the largest reductions in relative diversity occurring for the Native American and European ancestral segments (S11 Fig, S1 Supporting information).

**Fig 2.**
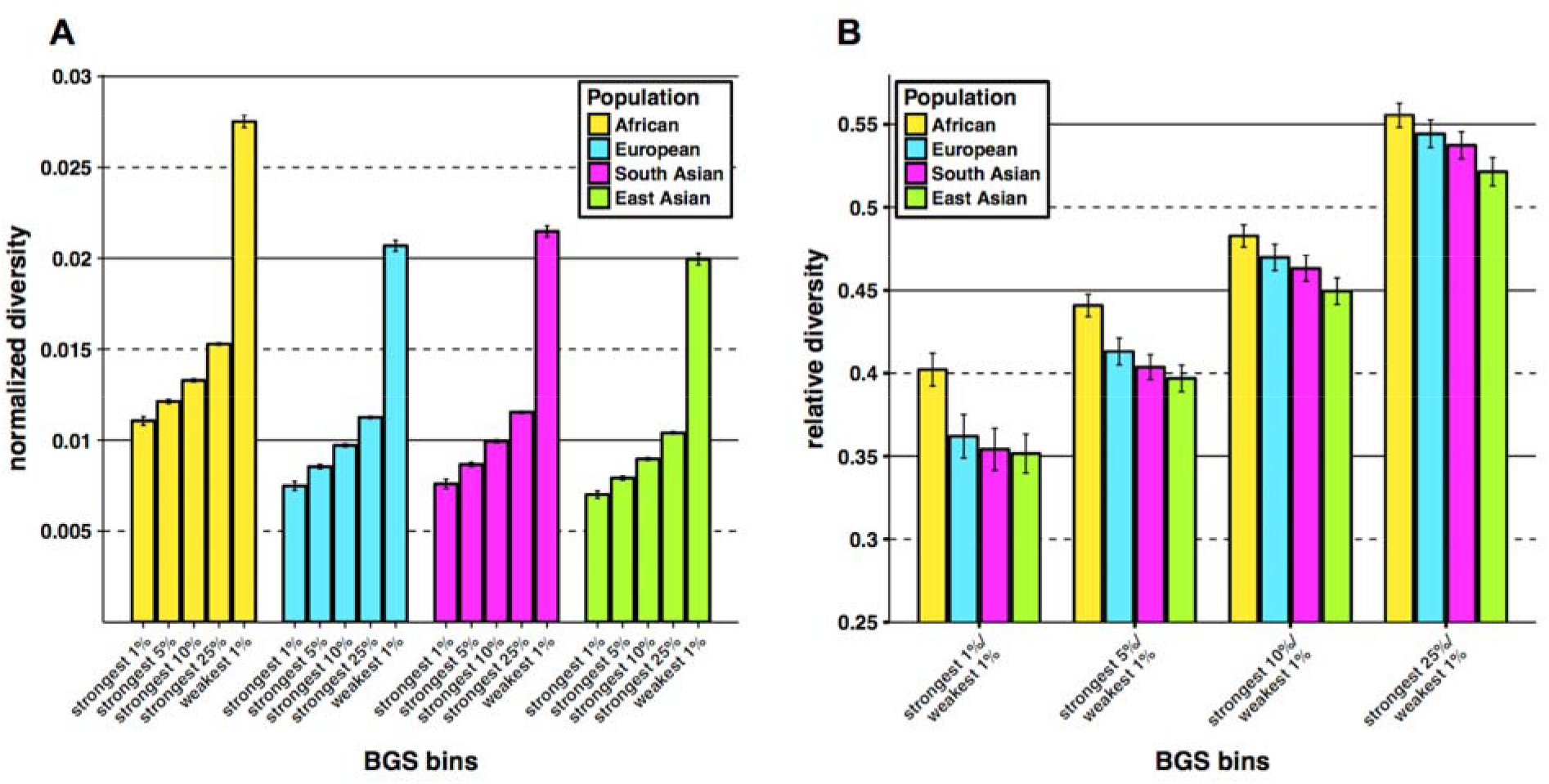
Normalized and relative diversity for Thousand Genomes Project (TGP) continental groups. (A) Normalized diversity (π/divergence) measured across the lowest 1%, 5%, 10% and 25% *B* quantile bins (strong BGS) and the highest 1% *B* quantile bin (weak BGS). (B) Relative diversity: the ratio of normalized diversity in the lowest *B* quantile bins (strong BGS) in (A) to normalized diversity in the highest 1% *B* quantile bin (weak BGS). Error bars represent ±1 SEM calculated from 1,000 bootstrapped datasets.

To test if demography has impacted selection at linked sites more recently in time, we also calculated the number of singletons observed per site (normalizing by divergence and using the same set of neutral filters as was used for the calculations of π) across the lowest and highest 1% *B* quantile bins. Singletons are, on average, the youngest variants within the genome and should better capture signals about more recent population history. We took the ratio of singletons observed per-site across these extreme *B* quantile bins to create a statistic called relative singleton density, which we term “ψ/ψ_0_.” We accounted for differences in population sample size by first projecting down all populations to 2N=170 (Materials and methods). Qualitatively, our measurements of ψ/ψ_0_ showed patterns in the opposite direction of our calculations of π/π_0_, with Africans exhibiting a lower ratio of ψ/ψ_0_ when compared to non-Africans (0.665-0.695 for Africans, 0.733-0.804 for non-Africans; Fig 3). These patterns suggest that the impact of demography on regions experiencing selection at linked sites is transient, with patterns of relative diversity between populations dependent on the time frame in which they are captured (see Discussion).

**Fig 3.**
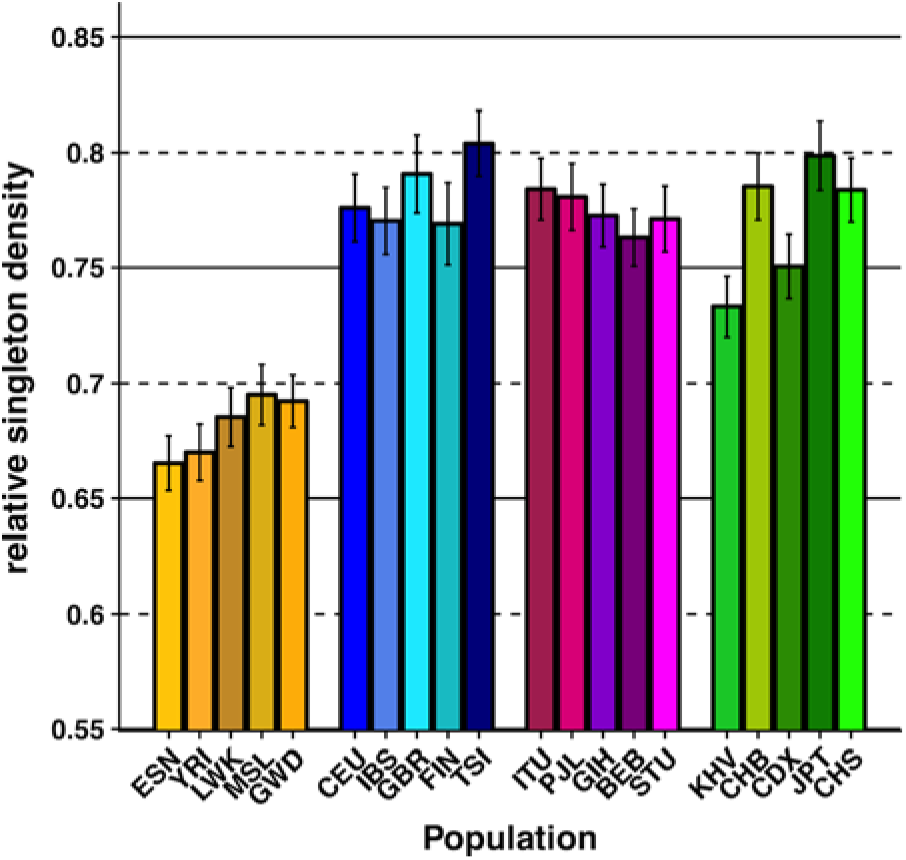
Relative singleton density for non-admixed populations of the Thousand Genomes Project (TGP) Relative singleton density measured by taking the ratio of singleton density in the lowest 1% *B* quantile bin to singleton density in the highest 1% *B* quantile bin (ψ/ψ_0_). Singleton density was normalized by divergence with Rhesus macaque. TGP population labels are indicated below each bar (see S12 Table in Supporting information for population label descriptions), with African populations colored by gold shades, European populations colored by blue shades, South Asian populations colored by violet shades, and East Asian populations colored by green shades. Error bars represent ±1 SEM calculated from 1,000 bootstrapped datasets.

### Selection at linked sites has shaped patterns of population differentiation

We have shown that selection at linked sites increases the rate of drift at neutral loci, and that this process is amplified by demographic changes like population bottlenecks. One might then expect that selection could also amplify population differentiation at linked neutral loci. This outcome is obvious in the context of hitchhiking (where linked neutral loci sweep to high frequency) but is also expected with BGS [61,62], since decreases in diversity can accelerate differentiation. Here we quantify the magnitude of the effect of BGS on population differentiation in humans, and find that population differentiation at neutral loci is indeed highly correlated with *B* (the inferred strength of BGS; Fig 4 and Table 1). Specifically, we divided the genome into 2% quantile bins based on the genome-wide distribution of *B* and measured *F*_ST_ in each bin for all pairs of populations from different continental groups [63]. We then performed simple linear regression using *B* as an explanatory variable and *F*_ST_ as our dependent variable with the linear model *F*_ST_ = *β_0_* + *β_1_B* + ε. We found that across all 150 population comparisons (i.e., the “Global” estimate in Table 1), *B* explained 26.9% of the change in *F*_ST_ across the most extreme *B* values, was robust to outliers [64] (S6 Table in Supporting information), and dominated the effects of local recombination rate (see Supporting information).

**Table 1.** Regression coefficient estimates for linear regression of *F_ST_* on 2% quantile bins of B.

|  | AFR vs.<br>EASN | AFR vs.<br>EUR | AFR vs.<br>SASN | EUR vs.<br>SASN | EUR vs.<br>EASN | SASN vs.<br>EASN | Global |
| --- | --- | --- | --- | --- | --- | --- | --- |
| $\beta_0$<br>± SEM<br>(p) | 0.2044<br>± 0.0039<br>($< 1e-04$) | 0.1716<br>± 0.0031<br>($< 1e-04$) | 0.1596<br>± 0.0029<br>($< 1e-04$) | 0.0455<br>± 0.0011<br>($< 1e-04$) | 0.1216<br>± 0.0029<br>($< 1e-04$) | 0.0903<br>± 0.0023<br>($< 1e-04$) | 0.1322<br>± 0.0019<br>($< 1e-04$) |
| $\beta_1$<br>± SEM<br>(p) | -0.0434<br>± 0.0046<br>($< 1e-04$) | -0.0358<br>± 0.0037<br>($< 1e-04$) | -0.0355<br>± 0.0034<br>($< 1e-04$) | -0.0098<br>± 0.0013<br>($< 1e-04$) | -0.0173<br>± 0.0035<br>($< 1e-04$) | -0.0261<br>± 0.0027<br>($< 1e-04$) | -0.0280<br>± 0.0022<br>($< 1e-04$) |
| $r$<br>± SEM | -0.8363<br>± 0.0295 | -0.7441<br>± 0.0362 | -0.7794<br>± 0.0332 | -0.3847<br>± 0.0414 | -0.6220<br>± 0.0785 | -0.5968<br>± 0.0348 | -0.1292<br>± 0.0098 |

The first two rows give the regression coefficients for the linear model $F_{ST} = \beta_0 + \beta_1 B + \varepsilon$ , where $B$ represents the mean background selection coefficient for the bin being tested and $F_{ST}$ is the estimated $F_{ST}$ for all population comparisons within a particular pair of continental groups (given in the column header). The final column, “Global”, gives the regression coefficients for the linear model applied to all pairwise population comparisons (150 total). The correlation coefficient, $r$ , between $B$ and $F_{ST}$ for each comparison is shown in the bottom row. Standard errors of the mean (SEM) for $\beta_0$ , $\beta_1$ , and $r$ were calculated from 1,000 bootstrap iterations (see Materials and Methods). P-values are derived from a two-sided t-test of the t-value for the corresponding regression coefficient.

**Fig 4.**
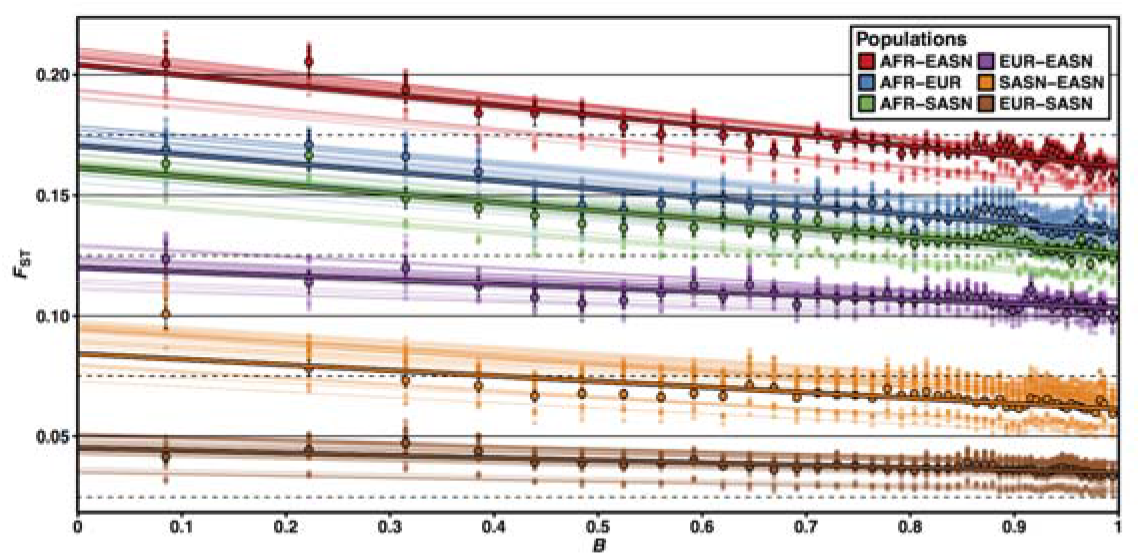
*F_ST_* is correlated with *B*. *F_ST_* measured across 2% quantile bins of B. Smaller transparent points and lines show the estimates and corresponding lines of best fit (using linear regression) for *F_ST_* between every pairwise population comparison for a particular pair of continental groups (25 pairwise comparisons each). Larger opaque points and lines are mean *F_ST_* estimates and lines of best fit across all Thousand Genomes Project (TGP) population comparisons between a particular pair of continental groups. Error bars represent ±1 SEM calculated from 1,000 bootstrapped datasets.

### Demographic inference in putatively neutral regions of the genome

One consequence of BGS and hitchhiking in driving patterns of neutral variation within and between human populations is that demographic inference could be substantially biased [51,52,65]. To assess the degree of bias in the context of human data, we fit a 13-parameter demographic model of African, European, and East Asian demography using only putatively neutral regions of the genome under the weakest effects of selection at linked sites (*B* ≥ 0.994) from a subset of TGP individuals with high coverage whole genome sequence data (see Materials and Methods). Our demographic model followed that of Gutenkunst et al. [7], with an ancient human expansion in Africa and a single out-of-Africa bottleneck followed by European- and East Asian-specific bottlenecks, as well as exponential growth in both non-African populations and migration between all populations. To make comparisons to previous studies that have used sequence data from coding regions or genes [7,22,23], which may be under strong BGS or hitchhiking effects, we also inferred demographic parameters using coding four-fold degenerate synonymous sites. Our inferred parameters for human demography were strikingly different between the two sets of sequence data (S1 Fig, S1 Table in Supporting information). Notably, inferred effective population size parameters were larger for contemporary population sizes when using four-fold degenerate synonymous sites versus ascertained neutral regions with *B* ≥ 0.994, with *N_e_* inferred to be 22%, 23%, and 29% larger for AFR, EUR, and EASN populations, respectively. This is despite the fact that the ancestral *N_e_* was inferred to be lower for four-fold degenerate synonymous sites *(N_e_* = 18,449 and 17,118, for neutral regions with *B* ≥ 0.994 and four-fold degenerate sites, respectively). This result may stem from the expected decrease in *N_e_* going into the past in regions of strong BGS, which can lead to inflated estimates of recent population growth [52] and has also been shown in simulation studies of synonymous sites under BGS [65]. Put more simply, the skew of the site-frequency spectrum towards rare variants in regions experiencing selection at linked sites mimics a population expansion, thus leading to erroneous inference.

### Simulations confirm that demographic effects can impact background selection

Using the demographic parameters inferred from neutral regions where *B* ≥ 0.994, we simulated patterns of neutral diversity with and without the effects BGS (see Materials and Methods). To measure the relative impact of BGS for each population, we then took the ratio of neutral diversity from BGS simulations (π) and neutral diversity from simulations without BGS (π_0_) to calculate relative diversity (π/π_0_). As expected, we found that BGS reduced relative diversity (π/π_0_ < 1) for all three populations in our simulations. However, non-African populations experienced a proportionally larger decrease in π/π_0_ compared to the African population (π/π_0_ = 0.43, 0.42, 0.41 in AFR, EUR, and EASN respectively). These results are comparable to (but not quite as extreme as) the effects we observed in the regions of the genome with the strongest effects of BGS for these population groups (Fig 1C). To understand how this dynamic process occurs, we sampled all simulated populations every 100 generations through time to observe the effect of population size change on π, π_0_, and the ratio π/π_0_ (Fig 5). We observed that there is a distinct drop in π and π_0_ at each population bottleneck experienced by non-Africans, with East Asians (who had a more severe bottleneck) experiencing a larger drop than Europeans. Fig 5C shows that the population bottlenecks experienced by non-African populations also reduces π/π_0_. Surprisingly, Africans also experienced a large drop in π/π0 (but less than non-Africans) even though they did not experience any bottlenecks. This was attributable to migration between non-Africans and Africans and this pattern disappeared when we ran simulations using an identical demographic model with BGS but without migration between populations (S7 Fig). This finding highlights an evolutionary role that deleterious alleles can play when they are transferred across populations through migration (see Discussion).

**Fig 5.**
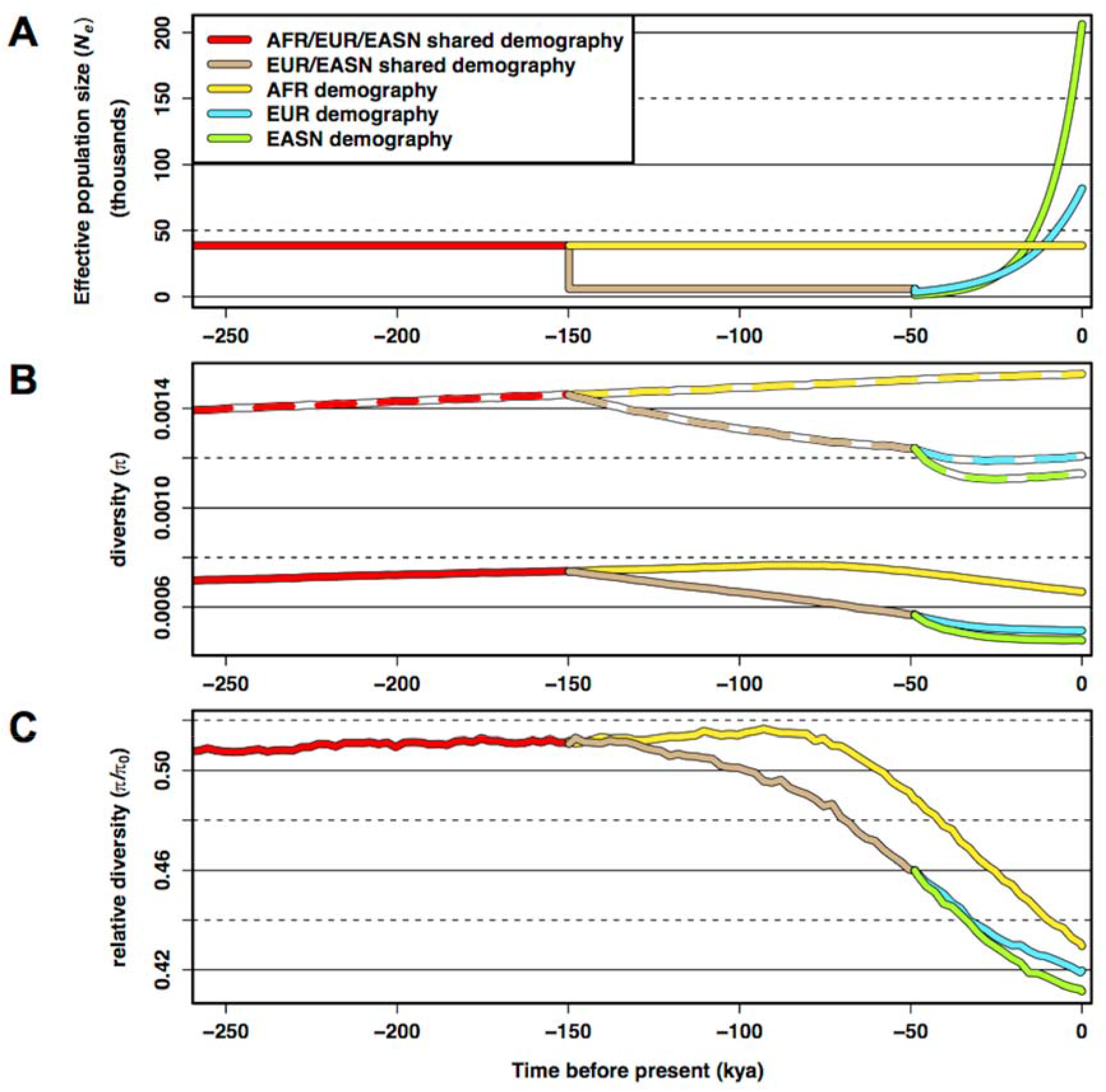
Simulations confirm that demographic events shape the impact of background selection (BGS) (A) Inferred demographic model from Complete Genomics Thousand Genomes Project (TGP) data showing population size changes for Africans (AFR), Europeans (EUR), and East Asians (EASN) as a function of time that was used for the simulations of BGS. (B) Simulated diversity at neutral sites across populations as a function of time under our inferred demographic model without BGS (π_0_ - dashed colored lines) and with BGS (π-solid colored lines). (C) Relative diversity (π/π_0_) measured by taking the ratio of diversity with BGS (π) to diversity without BGS (π_0_) at each time point. Note that the x-axes in all three figures are on the same scale. Time is scaled using a human generation time of 25 years per generation. Simulation data was sampled every 100 generations.

We also calculated ψ/ψ_0_ across these simulations and again qualitatively observed patterns similar to, but not as extreme, as our empirical calculations of ψ/ψ_0_. As with pairwise neutral diversity, BGS predictably decreased the density of singletons across all populations (S12 Fig A). However, Africans exhibited a lower ψ/ψ_0_ when compared to non-Africans (ψ/ψ_0_ = 0.79, 0.89, 0.91 in AFR, EUR, and EASN respectively). Calculating ψ and ψ_0_ through time showed that the population bottlenecks experienced by non-Africans led to strong decreases in both ψ and ψ_0_, with recent expansion in these populations then leading to large, rapid recoveries. Strong decreases in ψ/ψ_0_ after each population bottleneck were also observed, including a slight decrease in ψ/ψ_0_ in Africans that disappeared in the simulations without migration (S12 Fig B). While ψ/ψ0 for the European/East Asian ancestral population in the simulations with migration remained below that of Africans during the course of the Out-of-Africa bottleneck, we observed a rapid recovery in ψ/ψ_0_ for this population in the simulations without migration (compare bottoms panels, S12 Fig A and B). This suggests that for populations experiencing a sustained population bottleneck, the response of singletons to changes in the weakened intensity of BGS is quite rapid, especially when compared to patterns of π/π0 (compare Fig S7 C to Fig S12 B bottom panel). However, population migration mitigates this pattern. However, regardless of whether migration between populations was simulated, BGS seemed to have little effect on singleton density recovery in Europeans and Asians once population expansion occurred.

Our simulations were based on the functional density found in a 2 Mb region of the human genome with the lowest *B* values and, thus, where BGS was inferred to be strongest (chr3: 48,600,000-50,600,000). There, 20.46% of sites were either coding or conserved non-coding (see Materials and Methods). Thus, the fraction of the genome experiencing deleterious mutation in our simulations of strong BGS was 0.2046. The patterns we observed in these simulations likely represent an upper bound on the strength of BGS in the human genome. Since the strength of BGS is dependent upon the density of sites experiencing deleterious mutation within a given region (or more formally, *U*, which is the product of the per-site deleterious mutation rate and the number of sites experiencing deleterious mutation [66]), we simulated weaker effects of BGS by reducing the fraction of sites experiencing purifying selection (keeping the distribution of selective effects constant, see Materials and Methods). When the fraction of sites experiencing selection was decreased 2-4 fold in our simulations, we continued to observe a stepwise decrease in π/π_0_ while maintaining the specific rank order of African, followed by European, and then East Asian populations (S8 Fig). As expected, π/π_0_ increased for all populations as the fraction of sites that were simulated as deleterious decreased (π/π_0_ = 0.641 vs. 0.802, 0.62 vs. 0.777, and 0.611 vs. 0.777 for AFR, EUR, and EASN when the fraction of sites experiencing selection was reduced to 0.1023 and 0.05115, respectively). These simulations resulted in π/π0 values much larger than we observed in data (Figs 1C, 2B).

## Discussion

In our analyses of thousands of genomes from globally distributed human populations, we have confirmed that the processes of demography and selection at linked sites govern neutral variation across the genome. While this observation is not unexpected, we have characterized the dynamic consequence of non-equilibrium demographic processes in regions experiencing selection at linked sites in humans. We find that demography (particularly population bottlenecks) can amplify the consequences of selection at linked sites. To remove any possible biases that would influence our results, we controlled for functional effects of mutations, variability in mutation along the genome, potential sequencing artifacts, GC-based gene conversion, the potential mutagenic effects of recombination hotspots, and also removed regions of the genome with signatures of positive selection. None of these factors qualitatively affected our results.

We do recognize that one caveat of our controls is the fact that divergence itself is not independent of BGS [67], and this may present biases when using divergence to control for variation in mutation rate along the genome. This is because the rate of coalescence in the ancestral population of two groups will be faster in regions of strong BGS compared to regions of weak BGS due to the lower *N_e_* of the former, leading to a decrease in overall divergence in those regions. To limit the contribution of such biases in ancestral *N_e_* to divergence, we use rhesus macaque since it is more distantly related to humans than other primate species such as orangutan or chimpanzee (human-rhesus divergence: 29.6 MYA; [68]). Furthermore, if *B* truly captures the effects of BGS, then normalizing by the lower divergence that is characteristic of strong BGS bins and the higher divergence that is characteristic of weak BGS bins should make any differences between the two smaller, not greater. In fact, for our calculations of relative diversity in which we skip the normalization step, the differences in diversity between the lowest 1% and highest 1% *B* value bins are greater and give a lower ratio of relative diversity (π/π_0_ for AFR is 0.373 without the divergence step and 0.402 with the divergence step). A similar pattern is also observed for other continental groups (compare Fig 2 and S5 Fig). More importantly though, we should not expect the potential biases of our divergence step to contribute to the differences in relative diversity between each of the continental groups since biases in divergence estimates across the genome are based on the human reference sequence and are therefore identical across all human populations.

We also note that the estimates of *B* by McVicker et al. [6] may be biased by model assumptions concerning mutation rates and the specific sites subject to purifying selection, with the exact values of *B* unlikely to be precisely inferred. In fact, the *B* values provided by McVicker et al. range from 0 to 1, suggesting that some regions of the genome should be essentially devoid of diversity (but we do not observe this to be the case). Since our own analyses show that relative diversity has a lower bound at only ~0.35 in humans, the exact value of *B* itself should not be taken at face value. Rather, our primary motivation for using *B* was to ascertain regions that should be on the extreme ends of the genome-wide distribution of regions experiencing selection at linked sites, for which *B* should provide a good assessment. A study by Comeron et al. [32] that investigated BGS in *Drosophila* and utilized the same model of BGS as McVicker et al. found that biases presented by model assumptions or mis-inference on the exact value of *B* do not significantly change the overall rank order for the inferred strength of BGS across the genome. Thus we, expect McVicker et al.’s inference of *B* to provide good separation between the regions experiencing the weakest and strongest effects of selection at linked sites within the human genome, with model misspecification unlikely to change our empirical results.

While the effects of selection at linked sites in humans captured in our analyses could in principle include the consequences of positive selection, such as soft-sweeps and classic selective sweeps, we applied stringent filters to remove any such regions before our analysis (Materials and methods, S1 Appendix). We therefore expect that most of the resulting patterns we observe can be attributed to the consequences of BGS. Recently, a joint model of classic selective sweeps and BGS was applied to *Drosophila* and predicted that BGS has had a ~1.6 to 2.5-fold greater effect on neutral genetic diversity than classic selective sweeps [33], with another study giving similar results [69]. We should expect this magnitude to be even greater for humans, since classic selective sweeps were found to be rare in recent human evolution [41] and adaptive substitutions in the human genome are much less frequent than in *Drosophila* [5,70,71]. Nonetheless, we cannot rule out all contributions from hitchhiking to our results. In fact, our own simulations of BGS fail to capture the complete effects of linked selection on reducing π/π_0_ in different human populations (compare Figs 1C and 5C) and the additional contribution of hitchhiking to humans, which we did not simulate, may explain this discrepancy.

Non-equilibrium demography has also been of recent interest in regards to its impact on patterns of deleterious variation across human populations (often referred to as genetic load), with initial work showing that non-African populations have a greater proportion of non-synonymous deleterious variants that are homozygous compared to synonymous variants [56]. Similar results in human founder populations [57,72], *Arabadopsis* [73], and domesticated species such as dogs [12] and sunflowers [74] further demonstrate the pervasive impact that demography has on influencing the relative amount of deleterious variation across a variety of populations and species. Since BGS is a function of deleterious variation, it is not surprising that we also witness differences in π/π0 across human populations that have experienced different demographic histories. These effects are likley ubiquitous across other species as well. However, there has been recent contention about whether the previously described patterns of increased deleterious variation are driven by a decrease in the efficacy of natural selection (thus resulting in increased genetic load) or are solely artifacts of the response of deleterious variation to demographic change [58,75–78]. Recently, Koch et al. [55] investigated the temporal dynamics of demography on selected sites within humans and observed that after a population contraction, heterozygosity at selected sites can undershoot its expected value at equilibrium as low-frequency variants are lost at a quicker rate before the recovery of intermediate frequency variants can occur. In the context of both BGS and hitchhiking, which skew the site frequency spectrum of linked neutral mutations towards rare variants [26,66,79,80], we also expect a transient decrease in diversity as low-frequency variants are lost quickly during a population contraction. Indeed, as evident from our simulations of BGS and demography, immediatley after a population bottleneck, rapid losses in singleton density can occur, leading to transient decreases in ψ/ψ_0_. However, the recovery in singleton density is also quite rapid, while the recovery in π and π/π_0_ is quite slow. This is due to the fact that higher frequency variants, which contribute a greater amount to π, take a longer amount of time to recover after a population contraction compared to lower-frequency variants such as singletons. However, Koch et al. also demonstrated that the effect of demography on diversity is only temporary and that long-term diversity at selected sites approaches greater values once equilibrium is reached. We also stress these temporal effects on the patterns of relative neutral diversity that we observe here. In the context of BGS, if no population recovery occurs following a contracation, deleterious variants may behave more neutrally, weaking the overall effect of BGS and causing π/π_0_ to become larger than in the ancestral, pre-bottleneck population.

The temporal effects of non-equilbrium demographics on patterns of π/π_0_ and ψ/ψ_0_ may also explain the conflicting results obtained in a similar study of selection at linked sites in teosinte and its domesticated counterpart, maize [59]. In that study, the authors observed that π/π_0_ was higher in maize, which underwent a population bottleneck during domestication (no bottleneck event was inferred for the teosinte population) but that ψ/ψ_0_ was also lower. This result is contrary to what we observed qualitatively between non-African and African human populations. However, the demographic models that have been inferred for maize and humans are quite different. Maize is inferred to have had a recent, major domestication bottleneck that was essentially instantaneous and followed by rapid exponential growth [59]. In contrast, demographic models for non-African humans suggest a much more distant bottleneck that was sustained over a longer period of time, and only recently have non-African populations experienced rampant growth (coinciding with the advent of agriculture). Given the vastly different demographic models of the two species, it is therefore not surprising that patterns of variation would be different.

The greater contemporary *N_e_* of non-Africans could theoretically result in a greater efficacy of purifying selection and, consequently, a stronger efficacy of BGS, leading to their observed lower π/π_0_ relative to Africans. However, it is very unlikely that this explains the observed patterns of relative diversity that we see. The greater contemporary population size of non-Africans has transpired only in the very recent past, with accelerated growth in Europeans occurring within the last few hundred generations [23,81–84]. Rather, our simulations indicate that the response of π in regions under BGS is driven by the ancient, sustained population contractions humans have experienced, with reductions in π/π_0_ occurring concomitantly with the out-of-Africa bottleneck and European-East Asian split bottleneck events (Fig 5) and continuing even after the European and East Asian expansion events. In addition, our empirical measurements of ψ/ψ_0_ demonstrate that in recent time, BGS has had a lower effect in non-Africans compared to Africans. Our simulations further corroborate this result. The greater sensitivity of regions under BGS to population contractions and accelerated genetic drift is a more likely explanation. Broadly, our results show that contemporary patterns of neutral diversity cannot easily be attributable to contemporary forces of selection but instead may be exhibiting signatures that are still dominated by older demographic events. Interestingly though, our simulations reveal an additional factor that can influence the impact of BGS within populations - migration between populations. We observe that the exchange of deleterious variants from populations that have experienced extensive bottlenecks to populations with a more stable demography can magnify the strength of selection at linked sites. In particular, our simulations show that both π/π_0_ and ψ/ψ_0_ decrease in Africans despite the fact that they are inferred to have been constant in size in their recent evolutionary history (Fig 5B). These patterns disappear when migration is removed (Fig S7, S12 Fig B), however more work is needed to definitively test this.

While we describe here the differential effects of non-equilibrium demography on neutral diversity in regions under strong and weak BGS, it is worth mentioning that differences in the reduction of neutral diversity in the genome between different populations have also been investigated at the level of entire chromosomes. In particular, analyses of neutral diversity comparing autosomes to non-autosomes (i.e., sex chromosomes and the mitochondrial genome [mtDNA]) have been conducted. Interestingly, these studies have shown that population contractions have impacted the relative reduction of neutral diversity between non-autosomes and autosomes in a similar fashion to what we have observed between regions of strong BGS and weak BGS, with the greatest losses occurring in bottlenecked populations. This was demonstrated in humans [85] and later modeled and shown in other species [86], with the explanation that stronger genetic drift due to the lower *N_e_* of non-autosomes causes diversity to be lost more quickly in response to population size reductions. Recent work in humans has confirmed such predictions by showing that relative losses of neutral diversity in the non-autosomes are greatest for non-Africans [87–89]. These studies, plus others [90], have also shown that there is strong evidence for a more dominant effect of linked selection on the sex chromosomes relative to the autosomes in humans.

Since linked selection is a pervasive force in shaping patterns of diversity across the genomes in a range of biological species [1], it has been provided as an argument for why neutral diversity and estimates of *N_e_* are relatively constrained across species in spite of the large variance in census population sizes that exist [47,91]. However, since population bottlenecks are common among species and have an inordinate influence on *N_e_* [20], demography has also been argued as a major culprit for constrained diversity [2,91–93]. Yet, as we show in humans, it is likely that patterns of neutral diversity are in fact jointly impacted by both of these forces, magnifying one another to deplete levels of diversity beyond what is expected by either one independently. This may play an even larger role in higher *N_e_* species such as *Drosophila,* where the overall distribution of *B* was inferred to be even smaller (i.e., exhibiting stronger BGS) than in humans [32]. In our work, we also identify a potentially substantial role for migration from smaller populations that harbor more strongly deleterious alleles on patterns of linked neutral diversity in large populations. Together, these combined effects may help provide additional clues for the puzzling lack of disparity in genetic diversity among different species [94].

Finally, our results also have implications for medical genetics research, since selection may be acting on functional regions contributing to disease susceptibility. Since different populations will have experienced different demographic histories, the action of linked selection may result in disparate patterns of genetic variation (with elevated levels of drift) near causal loci. Recent work has already demonstrated that BGS’s consequence of lowering diversity impacts power for disease association tests [95]. Our results indicate that this impact may be even further exacerbated by demography in bottlenecked populations, leading to potentially larger discrepancies in power between different populations. Overall, this should encourage further scrutiny for tests and SNP panels optimized for one population since they may not be easily translatable to other populations. It should also further motivate investigators to simultaneously account for demography and linked selection when performing tests to uncover disease variants within the genome [95–97].

## Materials and methods

### Data

2,504 samples from 26 populations in phase 3 of the Thousand Genomes Project (TGP) [9] were downloaded from ftp://ftp.ncbi.nlm.nih.gov/1000genomes/. vcftools (v0.1.12a) [98] and custom python scripts were used to gather all bi-allelic SNP sites from the autosomes of the entire sample set.

A subset of TGP samples that were sequenced to high coverage (~45X) by Complete Genomics(CG) were downloaded from ftp://ftp.ncbi.nlm.nih.gov/1000genomes/. After filtering out related individuals via pedigree analyses, we analyzed 53 YRI, 64 CEU, and 62 CHS samples (S2 Table). The cgatools (v1.8.0) listvariants program was first used to gather all SNPs from the 179 samples using their CG ASM “Variations Files” (CG format version 2.2). Within each population, the number of reference and alternate allele counts for each SNP was then calculated using the cgatools testvariants program and custom python scripts. Only allele counts across high quality sites (i.e., those classified as VQHIGH variant quality by CG) were included. Low quality sites (i.e., those with VQLOW variant quality) were treated as missing data. Only autosomes were kept. Non-bi-allelic SNPs and sites vio lating Hardy-Weinberg equilibrium (HWE) (p-value < 0.05 with a Bonferroni correction for multiple SNP testing) were also removed.

We collected 13 whole-genome sequenced KhoeSan samples (sequence-coverage: 2.5–50X, see S3 Table in Supporting information) from 3 studies [99–101] and used the processed vcf files from each of those respective studies to gather all bi-allelic polymorphic SNPs (i.e., the union of variants across all vcf files). SNPs were only retained if they were polymorphic within the 13 samples (i.e., sites called as alternate only within the sample set were ignored).

### Filtering and ascertainment scheme

Positions in the genome were annotated for background selection by using the background selection coefficient, B, which was inferred by McVicker et al. [6] and downloaded from http://www.phrap.org/othersoftware.html. *B* was inferred by applying a classical model of BGS [60], which treats its effects as a simple reduction in *N_e_* at neutral sites as a function of their recombination distance from conserved loci, the strength of purifying selection at those conserved loci, and the deleterious mutation rate. *B* can be interpreted as the reduced fraction of neutral genetic diversity at a particular site along the genome that is caused by BGS, with a value of 0 indicating a near complete removal of neutral genetic diversity due to BGS and a *B* value of 1 indicating little to no effect of BGS on neutral genetic diversity (*B* = π/π_0_ = N_e_/N_0_). Positions for *B* were lifted over from hg18 to hg19 using the UCSC liftOver tool. Sites that failed to uniquely map from hg18 to hg19 or failed to uniquely map in the reciprocal direction were excluded. Sites lacking a *B* value were also ignored. We focused our analyses on those regions of the genome within the top 1%, 5%, 10%, and 25% of the genome-wide distribution of *B* and within the bottom 1% of the genome-wide distribution of B. These quantiles correspond to the *B* values 0.095, 0.317, 0.463, 0.691, and 0.994, respectively.

A set of 13 filters (referred to as the “13-filter set”) were used to limit errors from sequencing and misalignments with rhesus macaque and to remove regions potentially under the direct effects of natural selection and putative selective sweeps (we ignore the linked selection effects of background selection). These filters were applied to all samples in phase 3 TGP (all filters are in build hg19) for all sets of analyses (see S4 Table in Supporting information for the total number of Mb that passed the described filters below for each particular *B* quantile):

1. Coverage/exome: For phase 3 data, regions of the genome that were part of the high coverage exome were excluded (see ftp://ftp.ncbi.nlm.nih.gov/1000genomes/ftp/technical/reference/exome_pull_down_targets/20130108.exome.targets.bed.README). This was done to limit biases due to differing levels of coverage across the genome and to remove likely functional sites within the exome.
2. phyloP: Sites with phyloP [102] scores > 1.2 or < −1.2 were removed to limit the effects of natural selection due to conservation or accelerated evolution. Scores were downloaded from http://hgdownload.cse.ucsc.edu/goldenPath/hg19/phyloP46way/.
3. phastCons: Regions in the UCSC conservation 46-way track (table: phastCons46wayPlacental) [103] were removed to limit the effects of natural selection due to conservation.
4. CpG: CpG islands in the UCSC CpG islands track were removed because of their potential role in gene regulation and/or being conserved.
5. ENCODE blacklist: Regions with high signal artifacts from next-generation sequencing experiments discovered during the ENCODE project [104] were removed.
6. Accessible genome mask: Regions not accessible to next-generation sequencing using short reads, according to the phase 3 TGP “strict” criteria, were removed (downloaded from ftp://ftp.ncbi.nlm.nih.gov/1000genomes/ftp/release/20130502/supporting/accessible_genome_masks/StrictMask/).
7. Simple repeats: Regions in the UCSC simple repeats track were removed due to potential misalignments with outgroups and/or being under natural selection.
8. Gaps/centromeres/telomeres: Regions in the UCSC gap track were removed, including centromeres and telomeres.
9. Segmental duplications: Regions in the UCSC segmental dups track [105] were removed to limit potential effects of natural selection and/or misalignments with rhesus macaque.
10. Transposons: Active transposons (HERVK retrotransposons, the AluY subfamily of Alu elements, SVA elements, and L1Ta/L1pre-Ta LINEs) in the human genome were removed.
11. Recent positive selection: Regions inferred to be under hard and soft selective sweeps (using iHS and iHH12 regions from selscan v1.2.0 [106]; S1 Appendix) within each phase 3 population were removed.
12. Non-coding transcripts: Non-coding transcripts from the UCSC genes track were removed to limit potential effects of natural selection.
13. Synteny: Regions that did not share conserved synteny with rhesus macaque (rheMac2) from UCSC syntenic net filtering were removed (downloaded from http://hgdownload.soe.ucsc.edu/goldenPath/hg19/vsRheMac2/syntenicNet/). Additionally, an extra set of filters was applied, but only for those estimates of diversity that controlled for GC-biased gene conversion and recombination hotspots:
14. GC-biased gene conversion (gBGC): Regions in UCSC phastBias track [107] from UCSC genome browser were removed to limit regions inferred to be under strong GC-biased gene conversion.
15. Recombination hotspots: All sites within 1.5 kb (i.e., 3 kb windows) of sites with recombination rates ≥ 10 cM/Mb in the 1000G OMNI genetic maps for non-admixed populations (downloaded from ftp://ftp-trace.ncbi.nih.gov/1000genomes/ftp/technical/working/20130507_omni_recombination_rates/) and the HapMap II genetic map [108] were removed. 1.5 kb flanking regions surrounding the center of hotspots identified by Ref. [109] (downloaded from http://science.sciencemag.org/content/sci/suppl/2014/11/12/346.6211.1256442.DC1/1256442_DatafileS1.txt) were also removed, except for the cases in which the entire hotspot site was greater than 3 kb in length (in which case just the hotspot was removed).

To generate a set of four-fold degenerate synonymous sites, all polymorphic sites that we retained from the high-coverage Complete Genomic samples were annotated using the program ANNOVAR [110] with Gencode V19 annotations. ANNOVAR and Gencode V19 annotations were also used to gather an autosome-wide set of fourfold degenerate sites, resulting in 5,188,972 total sites.

### Demographic inference

The inference tool dadi (v1.6.3) [7] was used to fit, via maximum likelihood, the 3-population 13-parameter demographic model of Gutenkunst et al. [7] to the 179 YRI, CEU, and CHS samples from the high coverage CG dataset of TGP. This sample set consisted of 53 YRI (African), 64 CEU (European), and 62 CHS (East Asian) samples. The demographic model incorporates an ancient human expansion in Africa and a single out-of-Africa bottleneck followed by European- and East Asian-specific bottlenecks, as well as exponential growth in both non-African populations and migration between populations. During the inference procedure, each population was projected down to 106 chromosomes, corresponding to the maximum number of chromosomes available in the CG YRI population. Sites were polarized with chimpanzee to identify putative ancestral/derived alleles using the chain and netted alignments of hg19 with panTro4 (http://hgdownload.soe.ucsc.edu/goldenPath/hg19/vsPanTro4/axtNet/), and the correction for ancestral misidentification [111] option in dadi was used. The 13-filter set described previously was applied to the CG data set, and an additional filter keeping only the autosomal sites in the top 1% of *B (B* ≥ 0.994) was also applied in order to mitigate potential biases in inference due to BGS [52,65] or other forms of linked selection [51]. After site filtering and correction for ancestral misidentification, a total of 110,582 segregating sites were utilized by dadi for the inference procedure. For optimization, grid points of 120, 130, and 140 were used, and 15 independent optimization runs were conducted from different initial parameter points to ensure convergence upon a global optimum. An effective sequence length (L) of 7.15 Mb was calculated from the input sequence data after accounting for the fraction of total sites removed due to filtering. In addition to the 13-filter set, this filtering included sites violating HWE, sites without *B* value information, sites that did not have at least 106 sampled chromosomes in each population, sites with more than two alleles, sites that did not have tri-nucleotide information for the correction for ancestral misidentification step, and sites treated as missing data. For calculating the reference effective population size, a mutation rate (μ) of 1.66 x 10^−8^ (inferred from Ref. [112]) was used. Using the optimized *θ* from dadi after parameter fitting, the equation θ = *4N_e_μL* was solved for *N_e_* to generate the reference effective population size, from which all other population *N_e_*’s were calculated. This same procedure was also used to infer demographic parameters from four-fold degenerate synonymous sites across the same set of samples. After site filtering (note that *B* and the 13-filter set were not included in the filtering step for four-fold degenerate synonymous sites), 41,260 segregating sites were utilized by dadi for the inference procedure, and an effective sequence length of 2.37 Mb was used for calculating the reference effective population size.

### Simulations

Forward simulations incorporating the results from the demographic inference procedure described above and a model of background selection were conducted using SFS_CODE [113]. For the model of background selection, the recombination rate, *ρ* and the fraction of the genome experiencing deleterious mutation were calculated using the 2 Mb region of chr3: 48,600,000-50,600,000, which has been subject to the strongest amount of BGS in the human genome (mean *B* = 0.002). A population-scaled recombination rate (*ρ*) of 6.0443 x 10^−5^ was calculated for this region using the HapMap II GRCh37 genetic map [108]. For ascertaining the fraction of sites experiencing deleterious mutation, the number of non-coding “functional” sites in this region was first calculated by taking the union of all phastCons sites and phyloP sites with scores > 1.2 (indicating conservation) that did not intersect with any coding exons. This amount totaled to 270,348 base pairs. Additionally, the number of coding sites was calculated by summing all coding exons within this region from GENCODE v19, which totaled to 138,923 base pairs. From these totals, the total fraction of deleterious sites, 0.2046, was generated.

The background selection model was simulated using a middle 30 kb neutral region flanked by two 1 Mb regions under purifying selection. From the calculated fraction of deleterious sites described above, 20.46% of sites in the two 1 Mb flanking regions were simulated as being deleterious. The mutation rate in our simulations for the deleterious sites and for neutral sites were both set to 1.66 x 10^−8^ [112]. Two distributions of fitness effects were used for the deleterious sites, with 66.06% of deleterious sites using the gamma distribution of fitness effects inferred across conserved non-coding regions by Ref. [114] (β = 0.0415, α = 0.00515625) and 33.94% of deleterious sites using the gamma distribution of fitness effects inferred across coding regions by Ref. [5] (β = 0.184, α = 0.00040244). The relative number of non-coding “functional” sites and coding exons described above determined the relative number of sites receiving each distribution of fitness effects in our simulations. Gamma distribution parameters were scaled to the ancestral population size of the demographic models used in Refs. [5,114]. An example of the SFS_CODE command for our simulations is in S1 Supporting information. To simulate varying levels of background selection strength, different total fractions of our original calculated deleterious fraction of 0.2046 were used (i.e., 5%, 10%, 25%, 50%, and 100% of 0.2046). However, the same relative percentage of non-coding and coding sites and mutation rate were used. These different simulated fractions of deleterious sites resulted in a reduced total deleterious mutation rate, *U*, which is the product of the per-site deleterious mutation rate and the total number of sites experiencing deleterious mutation [66]. Thus, weaker effects of BGS were simulated. To simulate only the effects of demography without background selection, only the 30 kb neutral region was simulated. 2,000 independent simulations were conducted for each particular set of the deleterious site fraction (2,000 x 6 = 12,000 total). Simulations output population genetic information every 100 generations and also at each generation experiencing a population size change (22,117 total generations were simulated), from which mean pairwise nucleotide diversity (π) and singleton density (ψ) was calculated across the 2,000 simulations.

### Population-specific calculations of diversity and singleton density

Mean pairwise genetic diversity (π) and singleton density (ψ) was calculated as a function of the *B* quantile bins described in “Filtering and ascertainment scheme” for each of the 20 non-admixed populations in phase 3 TGP and, for π, across 4 broad populations that grouped the 20 non-admixed populations together by continent (African, European, South Asian, and East Asian, see S12 Table in Supporting information). Additionally, only regions of the genome passing the 13-filter set were used in the calculations of π and ψ (see S4 Table in Supporting information for total number of Mb used in diversity calculations for each *B* quantile). When calculating ψ for each non-admixed phase 3 TGP population, the site-frequency spectrum was first projected down to 2N = 170 samples (the number of chromosomes in MSL, the smallest phase 3 population sample) using a hypergeometric distribution [7] from each population’s full site-frequency spectrum. This allowed for unbiased comparisons of singleton density between all populations. For estimates of diversity controlling for gBGC or recombination hotspots, the additional corresponding filters described in “Filtering and ascertainment scheme” were also used. Only 100 kb regions of the genome with at least 10 kb of divergence information with Rhesus macaque were used in π and ψ calculations (see “Normalization of diversity and divergence calculations with Rhesus macaque” below).

### Normalization of diversity/singleton density and divergence calculations with Rhesus macaque

To calculate human divergence with Rhesus macaque, we downloaded the syntenic net alignments between hg19 and rheMac2 that were generated by blastz from http://hgdownload.cse.ucsc.edu/goldenpath/hg19/vsRheMac2/syntenicNet/. We binned the human genome into non-overlapping 100 kb bins and calculated divergence within each bin by taking the proportion of base pair differences between human and Rhesus macaque. Gaps between human and Rhesus macaque, positions lacking alignment information, and positions that did not pass the 13-filter set described in “Filtering and ascertainment scheme” were ignored in the divergence estimate. Additionally, a separate set of divergence estimates were also made using the additional set of filtering criteria that removed those regions under gBGC or in recombination hotspots and were used for normalizing diversity in those measurements that controlled for gBGC and hotspots.

When normalizing diversity and singleton density by divergence, only 100 kb bins that had at least 10 kb of divergence information were used (21,100 bins total for 13-filter set, 20,935 bins total for the 13-filter set plus the additional gBGC and hotspot filters). Bins with less than 10 kb of divergence information were ignored. To make estimates comparable, in those measurements of diversity that did not normalize by divergence, diversity was still calculated using the same set of 100 kb bins that had at least 10 kb for estimating divergence.

### Calculations of population differentiation (*F*_ST_) and linear regression

*F_ST_* calculations were performed as a function of *B* between every pair of non-admixed phase 3 TGP populations not belonging to the same continental group (150 pairs total). We followed the recommendations in Bhatia et al. [63] to limit biases in *F_ST_* due to 1) type of estimator used, 2) averaging over SNPs, and 3) SNP ascertainment. Specifically, we 1) used the Hudson-based *F_ST_* estimator [115], 2) used a ratio of averages for combining *F_ST_* estimated across different SNPs, and 3) ascertained SNPs based on being polymorphic in an outgroup (i.e., the KhoeSan). For ascertaining SNPs in the KhoeSan, we also performed filtering according to the filtering scheme described under “Filtering and ascertainment scheme.” For a position to be considered polymorphic in the KhoeSan, at least one alternate allele and one reference allele had to be called across the 13 genomes we utilized (see “Data”). These criteria left 3,497,105 total sites in the genome in the phase 3 dataset for *F_ST_* to be estimated across.

*F_ST_* was calculated across 2% quantile bins of *B* (based on the genome-wide distribution of B) for all pairwise comparisons of populations between a specific pair of continental groups (25 pairs total) or across all pairwise comparisons using all continental groups (150 pairs total). Simple linear regression was performed with the model *F_ST_* = *β_0_* + *β_1_B* + ε. The mean of the bounds defining each quantile bin was used when defining the explanatory variables for the regression. Linear regression, robust linear regression [64], and simple correlation were performed using the lm(), rlm(), and cor() functions, respectively, in the R programming language (www. r-project.org). To generate standard errors of the mean, this same procedure was performed on *F_ST_* results generated from each of 1,000 bootstrapped iterations of the data.

### Bootstrapping

#### Diversity Estimates

To control for the structure of linkage disequilibrium and correlation between SNPs along the genome, we partitioned the human genome into nonoverlapping 100 kb bins (these bins were identical to the 100 kb bins used for estimating divergence) and calculated mean pairwise diversity (π) or heterozygosity within each bin. We also normalized the diversity estimates by divergence within each bin. We then bootstrapped individual genomes by sampling, with replacement, the 100 kb bins until the number of sampled bins equaled the number of bins used for calculating the diversity point estimates (i.e., 21,100 bins or 20,935 bins total, depending on whether filters for gBGC and hotspots were applied). 1,000 total bootstrap iterations were completed and standard errors of the mean were calculated by taking the standard deviation from the resulting bootstrap distribution.

#### F_ST_

For bootstrapping *F_ST_*, the human genome was partitioned into non-overlapping 100 kb bins and were sampled with replacement until 28,823 bins were selected (the total number of non-overlapping 100 kb bins in the human autosomes). *F_ST_* was then calculated genome-wide for the bootstrapped genome as a function of *B* for every pairwise comparison of non-admixed phase 3 TGP populations not belonging to the same continental group. 1,000 total bootstrap iterations were completed and standard errors of the mean were calculated by taking the standard deviation from the *F_ST_* distribution calculated from all 1,000 iterations.

## Acknowledgements

We thank Lawrence Uricchio, Dominic Tong, Melissa Spear, Nicolas Strauli and three anonymous reviewers for helpful comments on the manuscript. The computations in this paper were run on the QB3 Shared Cluster at the University of California, San Francisco.

## Supporting information

S1 Appendix. Soft sweep detection and implementation in selscan v1.2.0.

S1 Supporting information.

**S1 Fig. Inference models inferred from TGP CG high *B* neutral regions and coding four-fold degenerate sites**. Solid lines are the inference results from running dadi on 53 YRI (African), 64 CEU (European), and 62 CHS (East Asian) TGP CG samples (projected down to 106 chromosomes during inference procedure) across neutral regions in the highest 1% *B* bin (*B* ≥ 0.994). Broken lines represent the inference results using the same CG samples, but with sequence data only from coding four-fold degenerate synonymous sites.

**S2 Fig. Diversity for TGP non-admixed populations while controlling for GC-biased gene conversion and recombination hotspots**. (A) Normalized diversity (π/divergence) measured across the lowest 1% *B* quantile bin (strong BGS). (B) Normalized diversity measured across the highest 1% *B* quantile bin (weak BGS). (C) Relative diversity: the ratio of normalized diversity for the lowest 1% *B* bin to normalized diversity for the highest 1% *B* bin (π/π_0_). Error bars represent ±1 SEM calculated from 1,000 bootstrapped datasets.

**S3 Fig. Diversity for TGP non-admixed populations without normalizing by divergence with Rhesus macaque**. (A) Diversity (π) measured across the lowest 1% *B* quantile bin (strong BGS). (B) Diversity measured across the highest 1% *B* quantile bin (weak BGS). (C) Relative diversity: the ratio of diversity for the lowest 1% *B* bin to diversity for the highest 1% *B* bin (π/π_0_). Error bars represent ±1 SEM calculated from 1,000 bootstrapped datasets.

**S4 Fig. Diversity for TGP continental groups while controlling for GC-biased gene conversion and recombination hotspots**. (A) Normalized diversity (π/divergence) measured across the lowest 1%, 5%, 10% and 25% *B* quantile bins (strong BGS) and the highest 1% *B* quantile bin (weak BGS). (B) Relative diversity (π/π_0_) for the lowest 1%, 5%, 10%, and 25% *B* bins. Error bars represent ±1 SEM calculated from 1,000 bootstrapped datasets.

**S5 Fig. Diversity for TGP continental groups without normalizing by divergence with Rhesus macaque**. (A) Diversity (π) measured across the lowest 1%, 5%, 10% and 25% *B* quantile bins (strong BGS) and the highest 1% *B* quantile bin (weak BGS). (B) Relative diversity (π/π_0_) for the lowest 1%, 5%, 10%, and 25% *B* bins. Error bars represent ±1 SEM calculated from 1,000 bootstrapped datasets.

**S6 Fig. *F*_ST_ measured across joint bins of *B* and recombination rate for different TGP continental groups.** The left panels of S6 Figs A-E show *F*_ST_ measured as a function of 25 4% recombination rate quantile bins conditional on three 2% *B* quantile bins (note log scale of x-axis for recombination rate). The right panels of S6 Figs A-E show *F*_ST_ measured as a function of 25 4% *B* quantile bins conditional on three 2% recombination rate quantile bins. The following continental group comparisons are shown for each plot: (A) African vs. European, (B) African vs. East Asian, (C) European vs. South Asian, (D) European vs. East Asian, (E) South Asian vs. East Asian. Smaller transparent points and lines show the *F*_ST_ estimates and corresponding lines of best fit (using linear regression) for each of the pairwise population comparisons within a particular pair of continental groups (25 comparisons total). Larger opaque points are mean *F*_ST_ estimates across all pairwise comparisons within a particular pair of continental groups (with bold lines showing their corresponding lines of best fit).

**S7 Fig. Simulations of diversity and relative diversity under BGS using a human demographic model without migration.** (A) Inferred demographic model from Complete Genomics TGP data. The demographic model used for the simulations in S7 Fig are identical to those used for Fig 5, except that migration parameters between all populations are set to 0. (B) Simulated diversity at neutral sites across populations as a function of time under our inferred demographic model without BGS (π_0_ - dashed colored lines) and with BGS (π - solid colored lines). (C) Relative diversity (π/π_0_) measured by taking the ratio of diversity with BGS (π) to diversity without BGS (π_0_) at each time point. Note that the x-axes in all three figures are on the same scale. Time is scaled using a human generation time of 25 years per generation. Simulation data was sampled every 100 generations.

**S8 Fig. Simulations of diversity and relative diversity under BGS using various fractions of sites experiencing deleterious mutation.** Values for the deleterious site fraction are provided in the header for each set of plots. Left column plots show results of simulations under a demographic model with migration between all human populations. Right column plots show results of simulations under a demographic model with no migration. Colored lines represent different populations though time and are identical to those in Fig 5 and S7 Fig The demographic model used is also identical to that in Fig 5 (for simulations with migration) and S7 Fig (for simulations without migration). Simulation data was sampled every 100 generations.

**S9 Fig. *F*_ST_ is not correlated with recombination rate.**

*F*_ST_ measured across 2% recombination rate quantile bins. The right panel of S9 Fig displays a narrower range of recombination rates to show detail. Smaller transparent points and lines show the estimates and corresponding lines of best fit (using linear regression) for *F*_ST_ between every pairwise population comparison for a particular pair of continental groups (25 pairwise comparisons each). Larger opaque points and lines are mean *F*_ST_ estimates and lines of best fit across all Thousand Genomes Project (TGP) population comparisons between a particular pair of continental groups. Error bars represent ±1 SEM calculated from 1,000 bootstrapped datasets.

**S10 Fig. *F*_ST_ between African (AFR) and South Asian (SASN) populations jointly across *B* and recombination rate.**

(A) *F*_ST_ as a function of 25 recombination rate bins (4% quantile bins) conditional on three different 2% *B* quantile bins (note log scale of x-axis for recombination rate). (B) F_ST_ as a function of 25 *B* bins (4% quantile bins) conditional on three different 2% recombination rate quantile bins. Smaller transparent points and lines show the Fst estimates and corresponding lines of best fit (using linear regression) for each of the pairwise comparisons of AFR vs. SASN Thousand Genomes Project (TGP) populations (25 comparisons total). Larger opaque points are mean *F*_ST_ estimates across all pairwise comparisons of AFR vs. SASN TGP populations (with bold lines showing their corresponding lines of best fit).

**S11 Fig. Comparing patterns of diversity between local ancestry segments of admixed samples and continental groups.**

(A) Normalized diversity (heterozygosity/divergence) and (B) Relative diversity: the ratio of normalized diversity in the lowest *B* quantile bins (strong BGS) in (A) to normalized diversity in the highest 1% *B* quantile bin (weak BGS). Local ancestry segments include African, European, and Native American ancestries. Continental groups include African, European, and East Asian populations. Error bars represent ±1 SEM calculated from 1,0 bootstrapped datasets.

**S12 Fig. Simulations of singleton density and relative singleton density.**

A) Results of simulations under a demographic model with migration between all human populations. B) Results of simulations under a demographic model with no migration. The second row of A) and B) shows measurements of singleton density (i.e., number of singletons observed per site) from simulations without BGS (ψ_0_ - dashed colored lines) and with BGS (ψ - solid colored lines). The bottom row of A) and B) shows corresponding relative singleton density (ψ/ψ_0_) measured by taking the ratio of singleton density with BGS (ψ) to singleton density without BGS (ψ0) at each sampled generation time point. The simulation data used for these measurements is identical to that of Fig 5 (for simulations with migration) and S7 Fig (for simulations without migration).

